# Molecular architecture of thylakoid membranes within intact spinach chloroplasts

**DOI:** 10.1101/2024.11.24.625035

**Authors:** Wojciech Wietrzynski, Lorenz Lamm, William H.J. Wood, Matina-Jasemi Loukeri, Lorna Malone, Tingying Peng, Matthew P. Johnson, Benjamin D. Engel

## Abstract

Thylakoid membranes coordinate the light reactions of photosynthesis across multiple scales, coupling the architecture of an elaborate membrane network to the spatial organization of individual protein complexes embedded within this network. Previously, we used *in situ* cryo-electron tomography (cryo-ET) to reveal the native thylakoid architecture of the green alga *Chlamydomonas reinhardtii* [1] and then map the molecular organization of these thylakoids with single-molecule precision [2]. However, it remains to be shown how generalizable this green algal blueprint is to the thylakoids of vascular plants, which possess distinct membrane architecture subdivided into grana stacks interconnected by non-stacked stromal lamellae. Here, we continue our cryo-ET investigation to reveal the molecular architecture of thylakoids within intact chloroplasts isolated from spinach (*Spinacia oleracea*). We visualize the fine ultrastructural details of grana membranes, as well as interactions between thylakoids and plastoglobules. We apply and further develop AI-based computational approaches for automated membrane segmentation and membrane protein picking [3], enabling us to quantify the organization of photosynthetic complexes within the plane of the thylakoid membrane and across adjacent stacked membranes. Our analysis reveals that, despite different 3D architecture, the molecular organization of thylakoid membranes in vascular plants and green algae is strikingly similar. In contrast to isolated plant thylakoids, where semi-crystalline arrays of photosystem II (PSII) appear to hold some membranes together, we find in intact chloroplasts that PSII is non-crystalline and has uniform concentration both within the membrane plane and across stacked grana membranes. Similar to *C. reinhardtii*, we observe strict lateral heterogeneity of PSII and PSI at the boundary between appressed and non-appressed thylakoid domains, with no evidence for a distinct grana margin region where these complexes have been proposed to intermix. Based on these measurements, we support a simple two-domain model for the molecular organization of thylakoid membranes in both green algae and plants.

## Introduction

Biological membranes and the proteins embedded within them are interwoven components of a dynamic multiscale system. Thylakoid membranes present one of the best examples of such a system, with interconnected relationships between membrane architecture, molecular organization, and physiological function [4, 5]. Therefore, to fully understand the light-dependent reactions of photosynthesis, it is crucial to describe thylakoid architecture with high precision in its most native state.

Thylakoids are the basic unit of oxygenic photosynthesis (with the exception of *Gloeobacteria*, which completely lack them [6, 7]). They form a variety of architectures in cyanobacteria and within the chloroplasts of plants and algae [8]. In vascular plants, thylakoids assemble into one of the most intricate membrane networks in living organisms: one continuous cisternae-like compartment that is folded and flattened into an interconnected network [9–11]. Thylakoid shape and packing within plant chloroplasts are optimized to maximize the surface area for light absorption and the coupled electron and proton transfer reactions conducted by photosynthetic electron transport chain complexes. The thylakoid network is divided into domains of appressed, stacked membranes (called grana) and non-appressed, free thylakoids (called stromal lamellae) that interconnect the grana. This architecture is crucial for the spatial separation of photosystem I (PSI) and photosystem II (PSII) and their associated light harvesting antenna complexes, LHCI and LHCII, thereby preventing the ‘spillover’ (and thus loss) of excitation energy from the short wavelength trap of PSII (680 nm) to the longer wavelength trap of PSI (700 nm) [12, 13]. The existence of two thylakoid domains and their dynamics has been shown to underpin a variety of regulatory mechanisms that control linear electron transfer (LET), cyclic electron transfer (CET) [14, 15], the PSII repair cycle [16], non-photochemical quenching (NPQ) [17], and the balance of excitation energy distributed to PSI and PSII [18, 19]. Despite over half a century of efforts using a multitude of techniques, including freeze-fracture electron microscopy (EM), negative-stain EM, atomic force microscopy, and fluorescence microscopy, our understanding of the organizational principles of thylakoid networks remains limited by resolution and the requirement for intactness of the system [19–25]. Indeed, numerous controversies remain, including the nature and organization of the grana margins and grana end membranes [26], the architecture of the connections between grana and stromal lamellae [9, 22, 27, 28], and the exact distribution of protein complexes such as ATP synthase (ATPsyn) and cytochrome *b*_6_*f* (cyt*b*_6_*f*) [29, 30].

A decade ago, we combined cryo-focused ion beam (FIB) milling [31, 32] with cryo-electron tomography (cryo-ET) [33, 34] to provide a detailed view of thylakoid architecture *in situ*, inside the chloroplasts of *Chlamydomonas reinhardtii* cells [1]. We then built upon this by leveraging advances in direct electron detector cameras and the contrast-enhancing Volta phase plate to understand the distribution of photosynthetic complexes within the native thylakoid network of *C. reinhardtii* cells [2]. Now, we apply *in situ* cryo-ET to the more challenging system of plant chloroplasts, utilizing an artificial intelligence (AI)-assisted approach [3, 35] to identify individual photosynthetic complexes and quantify their organization within networks of grana and stromal lamellae membranes.

## Results and Discussion

We isolated chloroplasts from six-week-old spinach plants then plunged them onto EM grids, thinned them by cryo-FIB milling, and finally imaged them by cryo-ET with defocus contrast (no Volta phase plate). To enhance contrast of the defocus imaging, we used AI-based denoising algorithms [36, 37]. The resulting tomograms provide a glimpse into the near-native organization of the spinach plastid (Figure 1A, Figure 1 – figure supplement 1, Figure 1 – video supplement 1). The double-membrane envelope encloses a protein-dense stroma (Figure 1B, segmentations: envelope in blue, stroma proteins in grey. Also note the high concentration of Rubisco throughout the chloroplast, e.g., Figure 2A, Figure 3; orange arrowheads point to Rubisco particles in Figure 3B). Suspended within the stroma, the thylakoid network is the main architectural feature of the chloroplast (Figure 1A-C, segmented in green). The thylakoids form a discrete network that is discontinuous from the surrounding chloroplast envelope. However, we did observe rare contact sites between the thylakoid and envelope membranes (Figure 1 – figure supplement 2), consistent with previous cryo-ET observations in *C. reinhardtii* [1, 38]. Spherical plastoglobules [39] decorated in putative protein densities were often seen interacting with thylakoid surfaces (Figure 1B-C, Figure 2, segmented in orange). We noticed holes through the thylakoid sheets precisely at the plastoglobule contact sites (Figure 2D-E). At those sites, plastoglobules attach to thylakoids along an extended stretch of high curvature thylakoid membrane, as opposed to making a single point contact to a flat thylakoid sheet (Figure 2F). This may help facilitate exchange between the two compartments. Segmentation of the membranes highlights the nature of the folded thylakoid network (Figure 1C-D, Figure 1 – figure supplement 3). The grana form well defined stacks of appressed membranes, which when viewed from the top, were revealed to be irregular cylinders with wavy edges (Figure 3A, Figure 3 – video supplement 1). Sometimes, thylakoid layers change length and extend laterally beyond the stack, breaking the grana’s vertical order (e.g., Figure 1 – figure supplement 3A). Distances between grana differ, and in some instances they almost merge with one another (Figure 5A-B, Figure 1 – figure supplement 3A). In our dataset, the number of thylakoids per grana stack varied from 4 to 22, with a mean of 10 (Figure 1E left). Every granum connects with the stromal lamellae membranes that spiral around each stack, in accordance with the helical staircase model (Figure 1C-D, Figure 3A) [9, 24]. Stromal lamellae extend from the stacks at variable angles, and after some distance, join other grana or rarely dead-end in the stroma (Figure 1C-D, Figure 1 – figure supplement 3A). Our cryo-ET volumes reveal the fine structural details of near-native membrane bilayers at high-resolution (e.g., Figure 4 left), allowing us to precisely quantify the morphometric parameters of thylakoids with sub-nanometer precision. We measured a membrane thickness of 5.1 ± 0. 3 nm, a stromal gap of 3.2 ± 0. 3 nm, a luminal thickness of 10.8 ± 2.0 nm, and a total thylakoid thickness (including two membranes plus the enclosed lumen) of 21.1 ± 1.8 nm (Figure 4) (for comparison see [1, 2, 30, 40, 41]).

**Figure 1.**
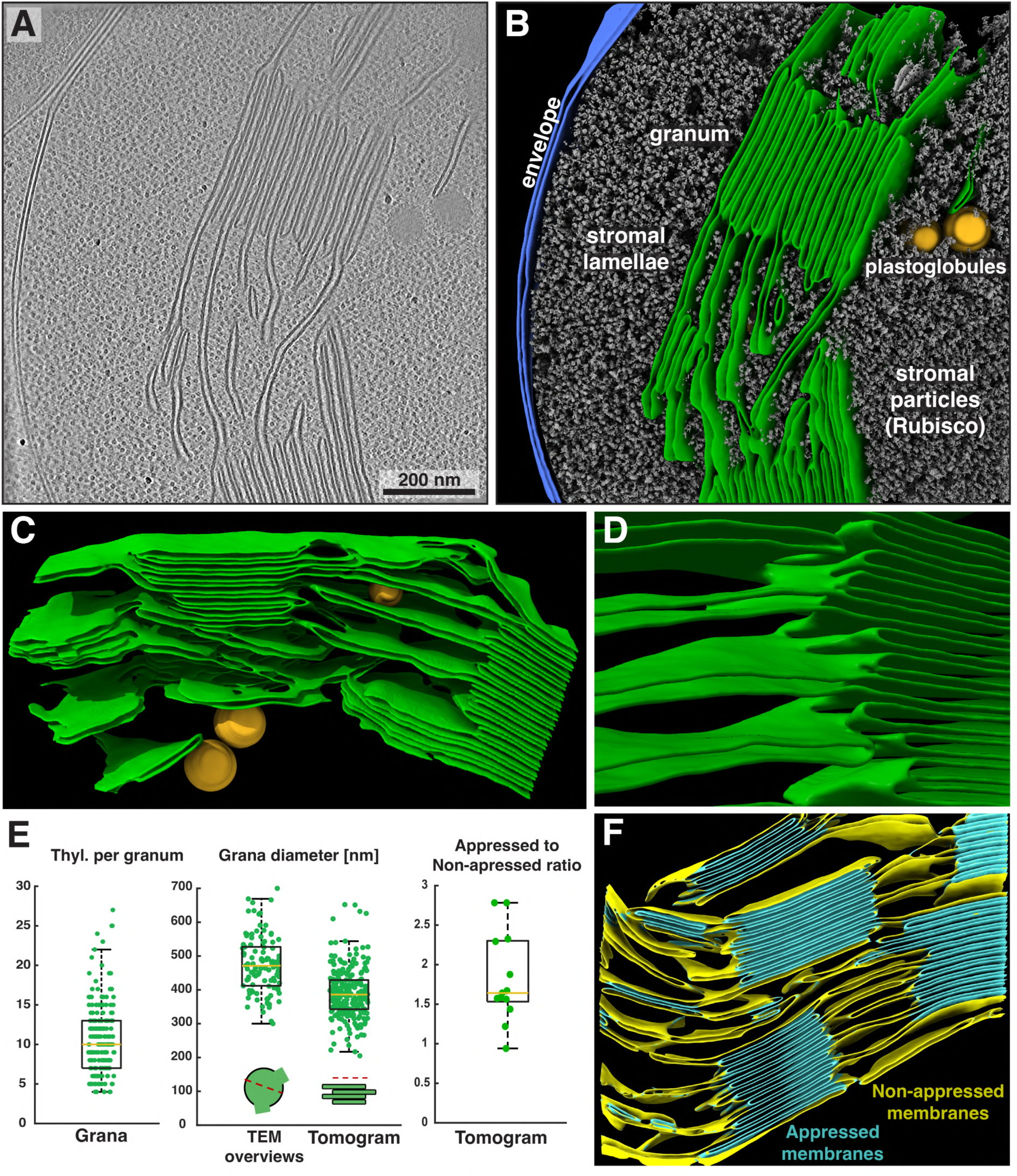
Overview of intact spinach chloroplasts visualized by cryo-ET. **A)** Tomographic slice through an intact spinach chloroplast. **B)** Corresponding 3D segmentation of the volume. Characteristic elements are labeled. Thylakoid membranes (green) are organized into stacks of appressed thylakoids (grana) and individual thylakoids connecting the grana (stromal lamellae). Note the intact chloroplast envelope (blue) and single thylakoid surrounded by two plastoglobules (orange). Stroma particles (grey) are a mix of different complexes but predominantly Rubisco. **C)** The same thylakoid network as in B, here shown from the opposite side. Note the inclined stromal lamellae spiraling around the granum on the left and a smaller plastoglobule sandwiched between the membranes. **D)** Close-up on the margin of a granum (from a different tomogram) showing the individual thylakoids transitioning into inclined stromal lamellae. In the first transition at the top of the image, note the hole in the thylakoid where the stromal lamella meets the granum. **E)** Box plots showing the number of thylakoids per granum (left), grana diameter measured in TEM overviews and in tomograms (middle), and the ratio of appressed to non-appressed membrane surfaces measured in selected tomograms. Box: 75 percentile, yellow lines: mean, whiskers: 95 percentile. **F)** Representative segmentation and classification of appressed (teal) and non-appressed (yellow) thylakoid membranes (quantification shown in E). See Figure 1 – video supplement 1 for an additional example of a chloroplast tomogram and segmentation of the thylakoid network.

**Figure 2.**
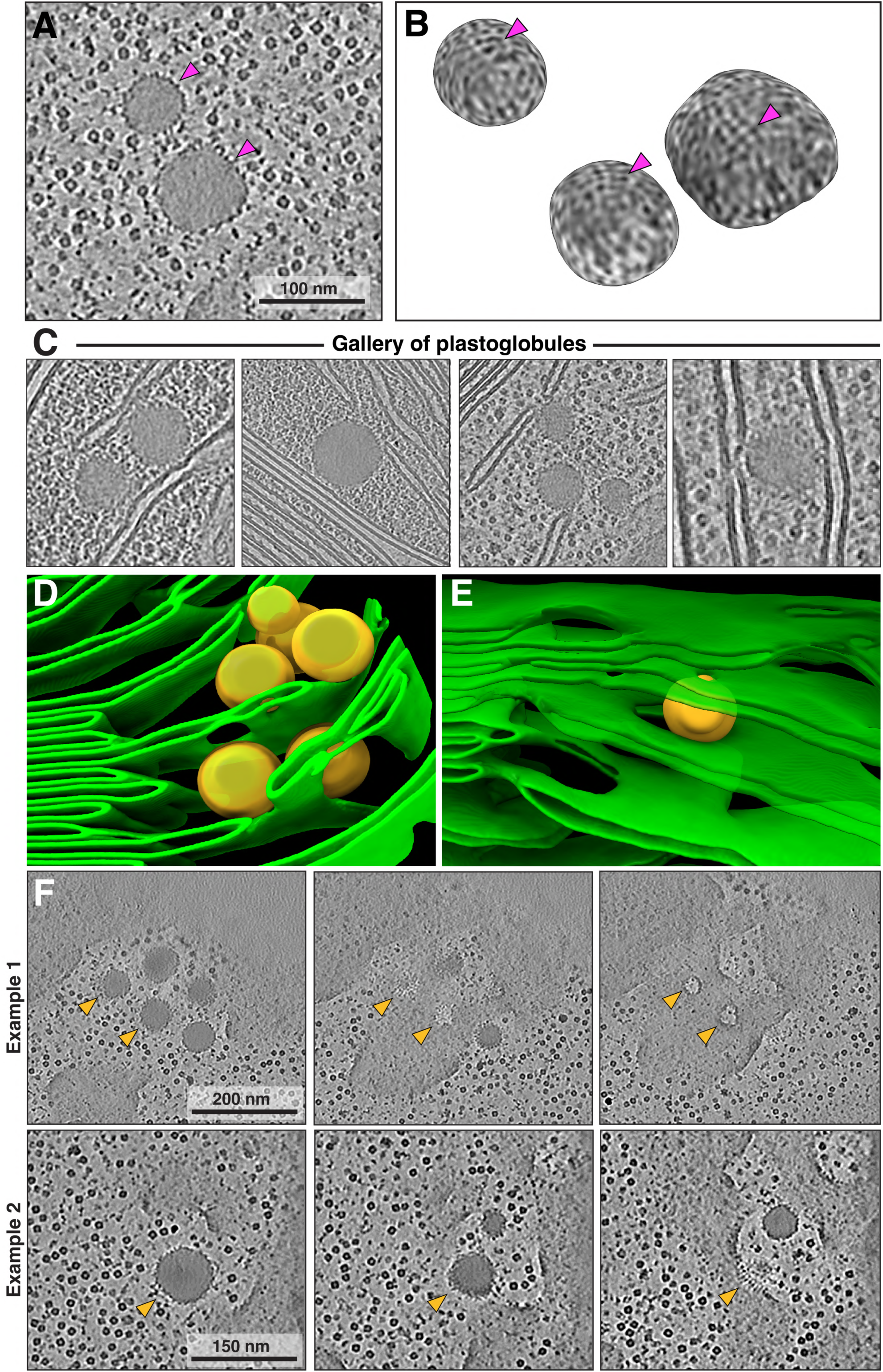
Fine details of plastoglobules interacting with thylakoids. **A)** Tomographic slice showing two plastoglobules: dense, spherical droplets that have surfaces covered with protein densities. **B)** 3D render of the surfaces of three plastoglobules. Note the ordered, small densities. Purple arrows in A and B indicate the protein coat. **C)** Gallery of tomographic slices showing plastoglobules in close proximity of thylakoids and the chloroplast envelope (second panel in the gallery of plastoglobules). **D-E)** Zoom-in on a segmentation of thylakoids with visible fenestrations in stromal lamellae. These holes may be caused by interactions with adjacent plastoglobules. **F)** Two examples of plastoglobules potentially interacting with thylakoids. Each example shows three consecutive tomographic z-slices. Note the protein-membrane interactions in example 2, with an array of small protein densities (potentially coming from the plastoglobule coat) mediating the contact between compartments.

**Figure 3.**
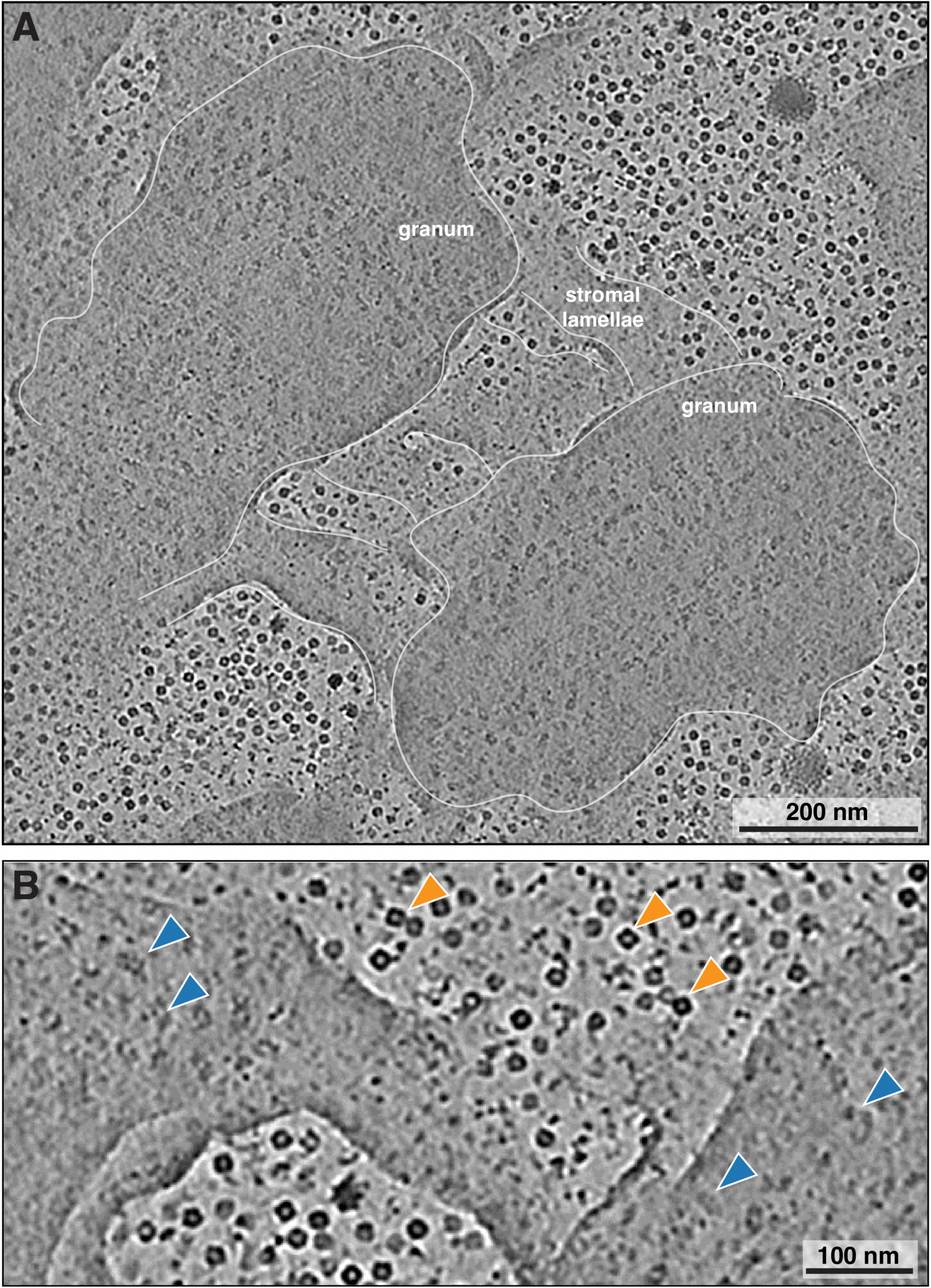
Top view of the interconnections between two grana. **A)** Averaged image of 40 tomographic slices (56 nm total averaged volume) from a tomogram denoised with the DeepDeWedge neural network [37] (high contrast, missing wedge inpainting), showing top views of two neighboring grana. Note the irregular shape of the grana, with wavy edges (traced with white line). Three stromal lamellae connect the two grana stacks. PSII particles are visible in the stacks. **B)** Close up of a region from A, showing stromal lamellae bridging two grana. Blue arrowheads: PSII particles in grana membranes; orange arrowheads: Rubisco particles In the stroma. See Figure 3 – video supplement 1 for tomographic slices through the entire volume, highlighting the dense membrane organization of the chloroplast.

**Figure 4.**
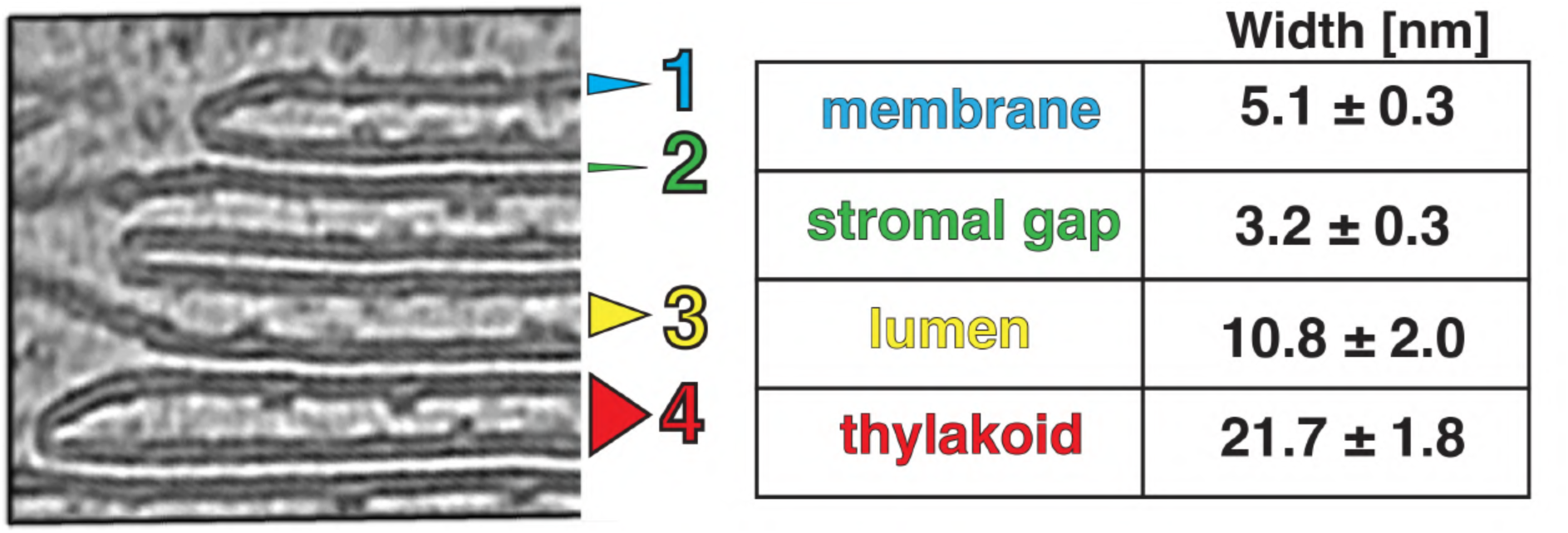
Thylakoid spacing and thickness measurements. Left: a zoom into thylakoids at the granum edge, showing the level of details observed in defocus cryo-tomograms denoised with cryo-CARE. Notice how the thylakoid tips stick to the appressed membrane, making them non-symmetrical. Right: quantification of thylakoid morphometric parameters. Membrane (1, blue) is the mean thickness of the lipid bilayer; stromal gap (2, green) is the mean distance between stacked thylakoids; lumen (3, yellow) is the mean thylakoid lumen width; thylakoid (4, red) is mean thickness of the thylakoid including two membrane bilayers plus the enclosed lumen (measured separately from the individual measurements of those features). Errors: SEM. Note the larger SEM values for lumen and thylakoid. This reflects the lumen variability between rigid appressed thylakoids of the granum and more labile stromal lamellae. Figure 4 – figure supplement 1 details the measurement procedure.

**Figure 5.**
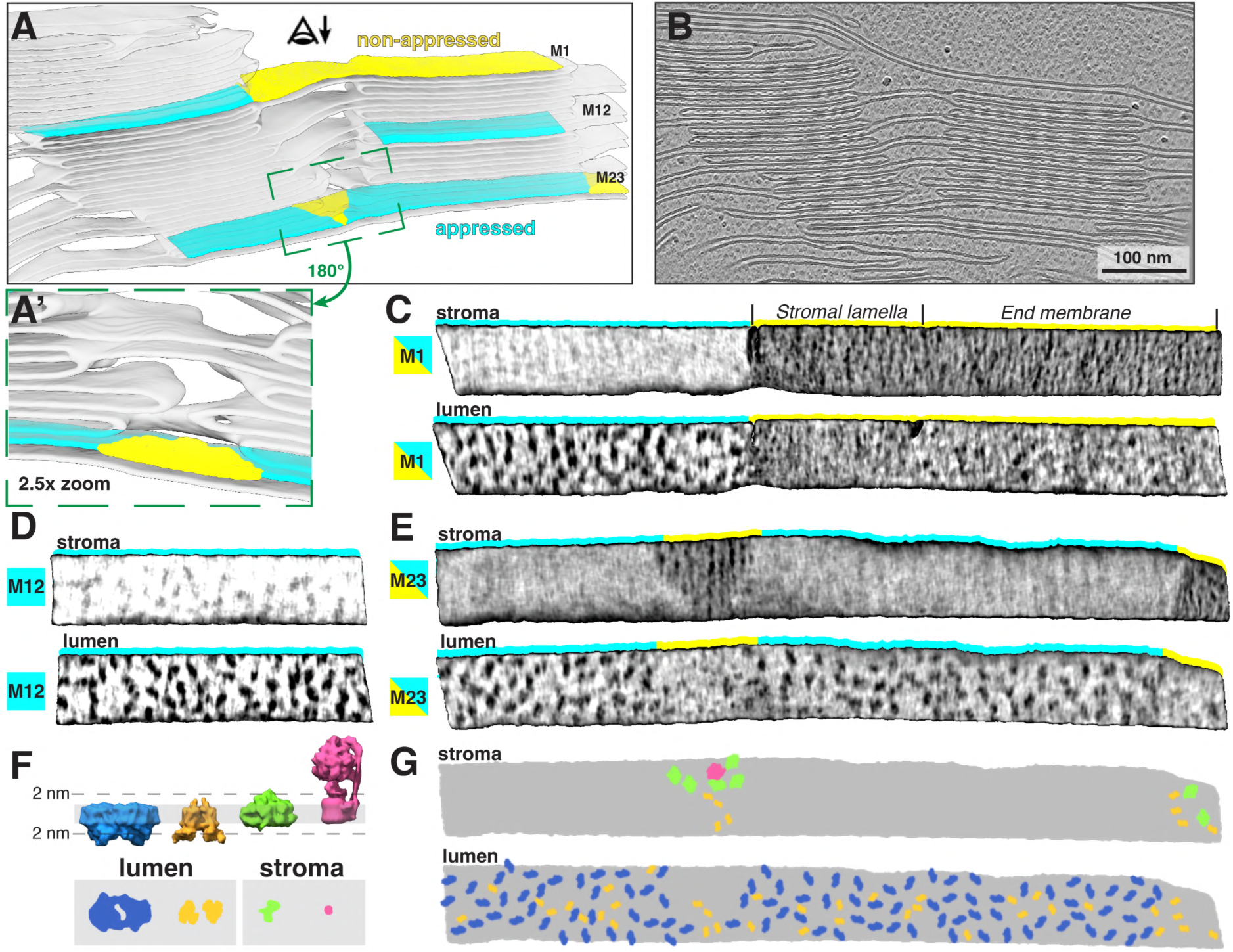
Lateral heterogeneity of PSII and PSI between appressed and non-appressed membranes. **A)** Segmentation of the thylakoid network showing two close grana. Three membrane pieces are highlighted: M1, an appressed membrane of the granum (teal) transitions into a stromal lamellae membrane (yellow) and then into an end membrane of the second granum (yellow); M12, an appressed membrane in the middle of the granum; M23, a membrane that spans two grana with a small unstacked stromal lamella in the middle (zoomed-in and rotated in green inset panel A’) and at the right end. The eye with the arrow indicates the viewing direction in membranograms. **B)** Slice through the corresponding tomographic volume. **C-E)** Membranograms showing both the luminal and stromal sides of the three membranes highlighted in A. Color strips indicate the membrane domain (blue: appressed, yellow: non-appressed). Stromal lamellae and end membrane are annotated above membranogram. All membranograms show densities 2 nm above the membrane surface (see F for reference). There are sharp boundaries between densities in the appressed and non-appressed regions. PSII densities are seen only on the luminal side of the appressed membranes, whereas PSI densities are seen only on the stromal side of non-appressed membranes. **F)** Top: Diagram showing side views of structures (blue: PSII, orange: cyt*b*_6_*f*, green: PSI, pink: ATPsyn) in a thylakoid membrane (grey). Bottom: membrane surface views showing densities that protrude from each structure 2 nm into the thylakoid lumen or stroma. **G)** Cartoon representation of the stromal and luminal surfaces of the membrane in E, with particle footprints pasted in (colored as in F).

It is important to note that individual tomograms represent only a small slice of the chloroplast volume (approximately 1200 x 1200 x 120 nm). This makes sampling of the larger micrometer-scale features less statistically relevant (see [42] and references within for alternative approaches). For example, grana diameter measured in our cryo-ET volumes is inherently underestimated due to FIB milling yielding only a narrow slice through the grana stack (Figure 1E middle). We therefore complemented these measurements with TEM overviews of FIB-milled lamellae, where many top views of grana were visible (Figure 1 – figure supplement 1B), facilitating more accurate diameter measurements (Figure 1E middle). Similarly, we are able to report the exact ratio between appressed and non-appressed membrane surface areas in our dataset (Figure 1E right, F), but we are aware that this ratio may not be representative of the entire chloroplast volume (see Limitations and Future Perspectives).

Thylakoid membranes are populated primarily by the photosynthetic complexes. Their distribution follows the principle of lateral heterogeneity [43]: PSII complexes are localized to appressed membranes, whereas PSI and ATPsyn complexes reside in the non-appressed regions [44–47]. The size of the stroma-exposed domains of PSI and ATPsyn (∼3.5 nm and ∼12 nm, respectively) prevents them from entering the narrow stromal gap between appressed thylakoids (∼3.2 nm, Figure 4). We were able to visualize particle distributions in appressed and non-appressed regions by projecting the tomogram volumes onto both the stromal and luminal surfaces of segmented membranes (Figure 5A, Figure 5 – figure supplement 1) to generate “membranograms” (Figure 5C-E, Figure 6A). Similar to *C. reinhardtii* [2] and as observed in other systems [28, 48, 49], large densities corresponding to the protruding oxygen evolving complexes (OEC) of PSII are visible on the luminal side of the appressed membranes (e.g., Figure 3, Figure 5C-E, Figure 3 – figure supplement 1D). These OEC densities mark the center positions of large membrane-embedded supercomplexes of PSII bound to trimers of LHCII [8, 50, 51], which do not extend from the membrane and thus are not visible in the membranograms. Interspersed between the PSII densities, we observe smaller dimeric particles that are consistent with the size of the lumen-exposed domains of cyt*b*_6_*f*, confirming the long-debated presence of this complex in appressed membranes of vascular plant thylakoids (e.g., [52]). In contrast, the stromal surface of the appressed membranes is generally smooth and featureless, consistent with the exclusion of PSI and ATPsyn, as well as the lack of large protrusions on the stromal faces of PSII, LHCII or cyt*b_6_f*. The PSII complexes abruptly disappear from the luminal face at the junction between appressed and non-appressed thylakoids (teal-to-yellow transitions in Figure 5C, E; Figure 5 – figure supplement 1A). Instead, both the luminal and stromal surfaces of the non-appressed thylakoids are covered by high number of smaller densities (Figure 5C, Figure 5 – figure supplement 1). The identity of these densities is difficult to assign with confidence, but a majority of them likely correspond to PSI (on the stromal side) and cyt*b*_6_*f* (on the luminal side) (see diagram in Figure 5F).

**Figure 6.**
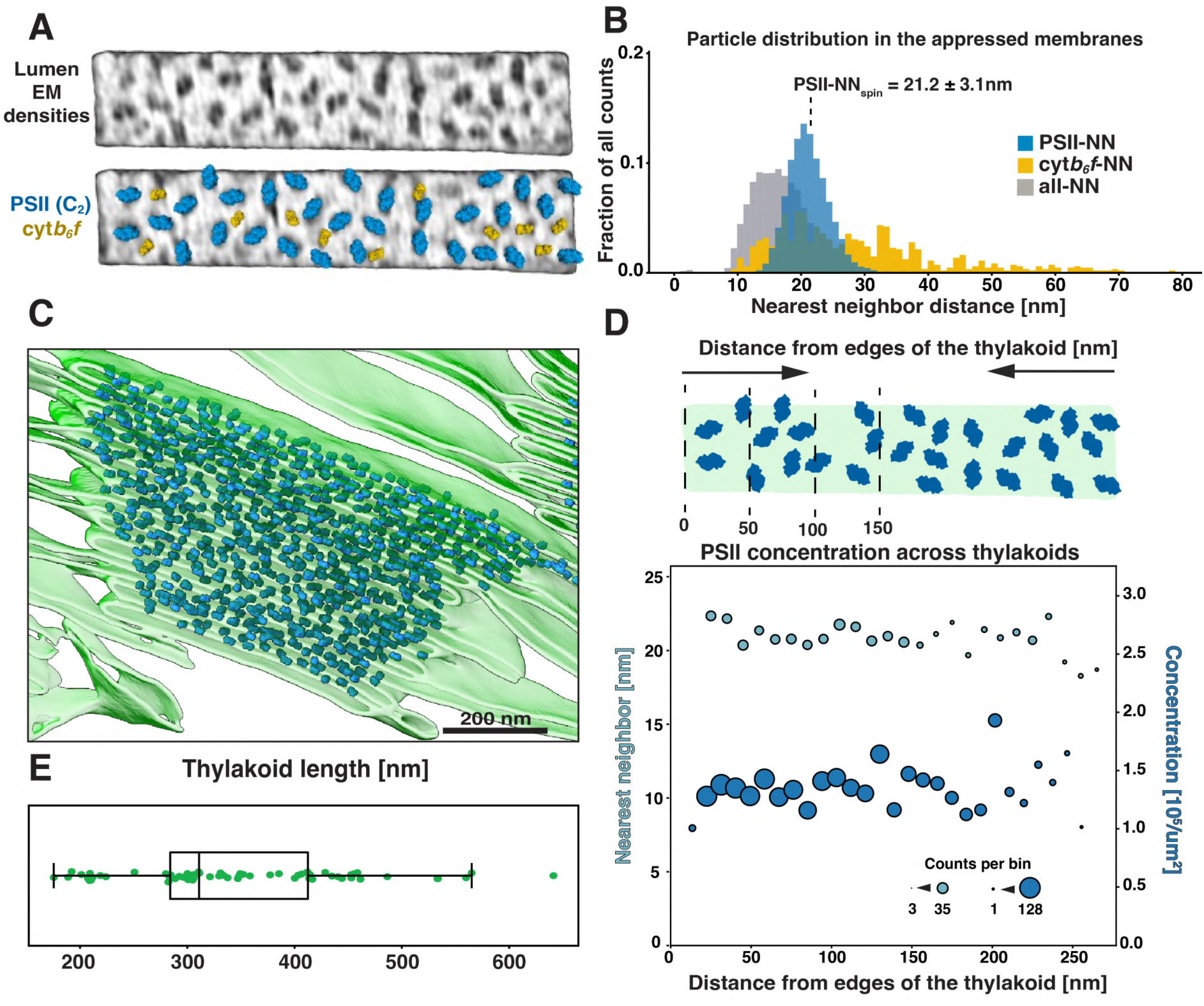
Distribution and concentration of PSII in grana membranes. **A)** Top: membranogram view of an appressed membrane piece with characteristic densities of dimers of PSII and cyt*b6f*. Bottom: same membrane with models of PSII and cyt*b6f* complexes fitted into the EM densities. **B)** Nearest neighbor distance plot of all the particles from the appressed membranes. Blue: PSII, orange: cyt*b6f*, grey: all particles. Note that “all” also includes unassigned particles of unknown identity. The dotted line labeled PSII-NNSpin indicates the mean (± SEM) center-to-center distance between neighboring PSII cores. Nearest neighbor distances are much more variable between two cyt*b6f* than between two PSII complexes. Identification of cyt*b6f* particles was possible only in the best resolved membranes; therefore, particle number and distribution estimation are not as reliable. **B)** Segmentation of a granum with all detected PSII particles pasted in. Note that we did not analyze PSII distribution at the very tips of thylakoids; this is why the tips look unoccupied. **D)** Plot showing PSII nearest neighbor center-to-center distances (light blue) and concentration (dark blue) as a function of the particle position from the edge of the granum, in 10 nm bins (diagrammed above the plot). Dot size represents number of counts in each bin. **E)** Plot of the lengths of grana thylakoids selected for the analysis in D. Thylakoids of significantly different lengths were selected to minimize any potential effect of the membrane size on the PSII distribution.

Biochemical isolation experiments often sub-divide the non-appressed membranes into stromal lamellae, grana margins, and grana end membranes on the basis of detergent solubilization and mechanical fragmentation, and have reported different protein compositions for the regions (for references see e.g., [26, 53]). In contrast, we observe little difference in the appearance and distribution of the densities on non-appressed membranes irrespective of whether they are immediately adjacent to grana (margins), extend away from them (stromal lamellae), or cap the grana stacks (grana end membranes) (Figure 5C, E; Figure 5 – figure supplement 1). Rather, we observe a simpler two-domain organization of photosynthetic complexes segregated into appressed and non-appressed regions, which corresponds to the original lateral heterogeneity model put forward by Andersson and Anderson [43, 54, 55]. It should be noted that the edges (tips) of the appressed membranes are difficult to visualize with membranograms, although their highly curved nature makes it unlikely that they could accommodate large photosynthetic complexes (see zoomed-in views of thylakoid tips in Figure 1D, Figure 4, Figure 1 – figure supplement 3B-D).

Using AI-assisted approaches [3, 35] confirmed by manual inspection, we detected essentially all particles populating appressed spinach membranes ([All_Spin_] = 2160 particles/µm^2^). Dimeric PSII complexes were clearly identified in the tomograms ([PSII_Spin_] = 1415 particles/µm^2^), enabling us to assign their center positions and orientations (Figure 6A, C) [56]. In the best quality tomograms, we were also able to manually assign dimeric cyt*b*_6_*f* particles in the appressed regions ([cyt*b*_6_*f*_Spin_] = 446 particles/µm^2^) (Figure 6A). Note that the total particle concentration, but not the PSII and cyt*b*_6_*f* subclasses, includes unknown densities as well as clipped particles at the edges of membrane patches selected for analysis. The PSII concentration and mean center-to-center distance between nearest neighbor complexes (PSII-NN_Spin_ = 21.2 ± 3.1 nm) (Figure 6B) is similar to what was reported in *Arabidopsis thaliana* by freeze fracture EM [57]. In contrast, our previous cryo-ET measurements of *C. reinhardtii* showed a slightly lower PSII concentration and correspondingly longer nearest neighbor distances within appressed thylakoid regions ([PSII_Chlamy_] = 1122 PSII/µm^2^, PSII-NN_Chlamy_ = 24.4 ± 5.2 nm)[2]. This discrepancy is in agreement with the relative sizes of the PSII-light harvesting complex II (PSII-LHCII) supercomplexes isolated from the respective species, with C_2_S_2_M_2_L_2_ supercomplexes from *C. reinhardtii* having a larger footprint in the membrane than C_2_S_2_M_2_ supercomplexes from vascular plants [58–60]. The concentration of PSII complexes is consistent between membranes (Figure 6B) and is independent of distance to the edge of the granum and thylakoid position in the stack (Figure 6E shows lengths of thylakoids analyzed in Figure 6D; we sampled membranes with variable lengths to depict the entire range of thylakoid sizes).

Using the observable orientations of OEC densities within the membrane plane, we placed footprints into the segmented membranes corresponding to structures of the C_2_ dimeric PSII core complex, the C_2_S_2_ PSII-LHCII supercomplex (isolated from spinach [59], PDB: 3JCU), and the C_2_S_2_M_2_ PSII-LHCII supercomplex (isolated from *A. thaliana* [60], PDB: 7OUI). When considering the PSII cores alone (C_2_), only rare events of complexes touching were observed (Figure 7A top row, D). Placing in the C_2_S_2_ supercomplexes resulted in a small in-plane clash between some particles (∼2.1% overlap of total footprint area, Figure 7A second row, D). The steric hindrance increased to ∼8.2% overlap when larger C_2_S_2_M_2_ supercomplexes were used (Figure 7A third row, D). Depending on which footprint was mapped in, the total membrane surface coverage by the PSII cores and the PSII-LHCII supercomplexes ranged from ∼20% to ∼55% (Figure 7B), leaving ample space in the membrane for additional free LHCII trimers and cyt*b*_6_*f*. Our observation of some clash between C_2_S_2_M_2_ supercomplexes suggests that different supercomplex variants probably coexist within the same membrane. Cumulative coverage of C_2_, C_2_S_2_, and C_2_S_2_M_2_ footprints when overlaying sequential membranes across the granum (known as the “granum crossection”) resulted in complete cross-section coverage after minimum of 6 membranes (3 consecutive thylakoids) in the case of the biggest supercomplex (Figure 7C).

**Figure 7.**
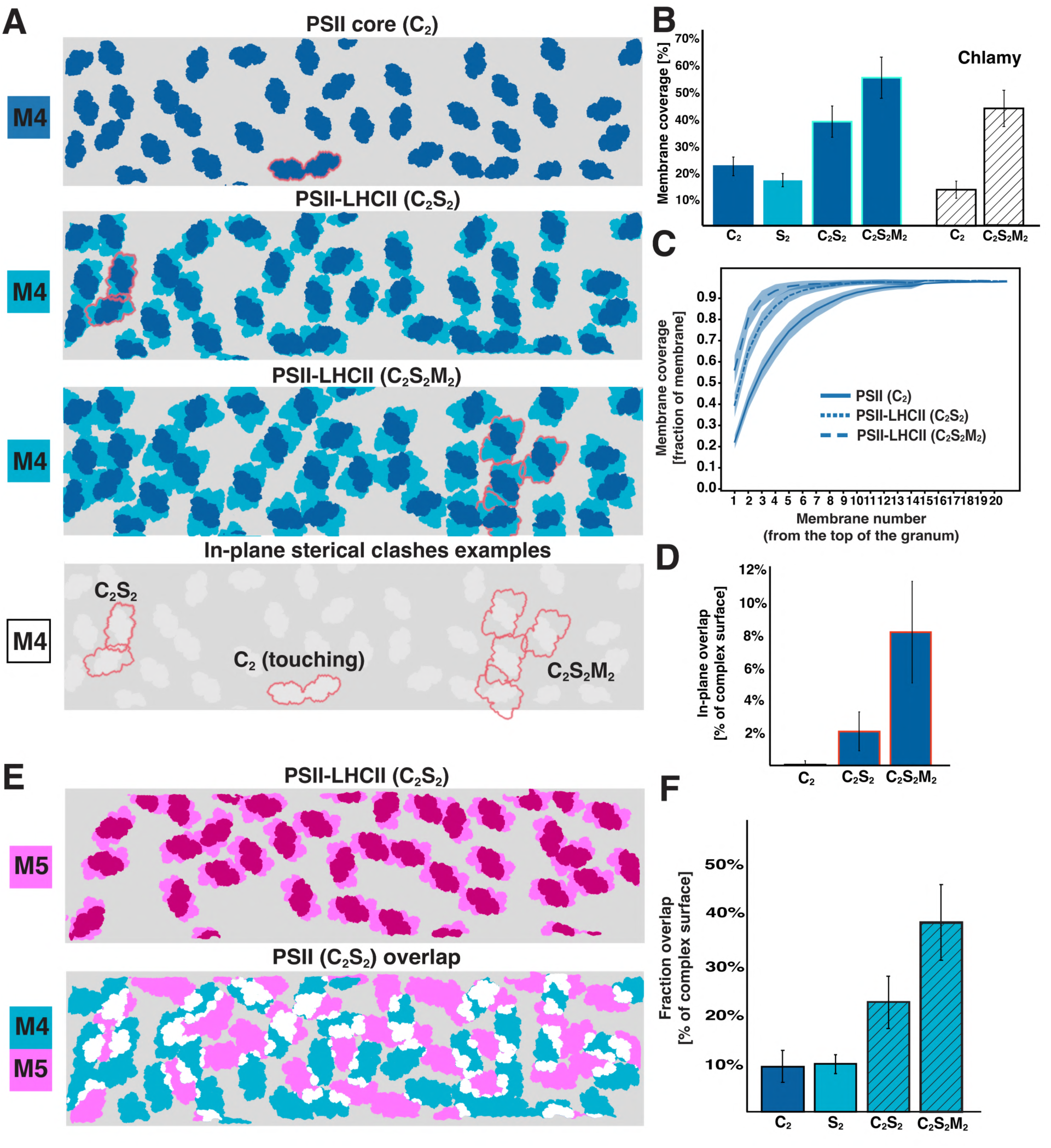
Occupancy of PSII and PSII-LHCII supercomplexes in grana membranes. **A)** Top panel: cartoon representation of an analyzed membrane patch (M4, fourth membrane in a granum) with footprints of all PSII core complexes (C2) placed into their respective positions. Middle panels: the same membrane patch with footprints of C2S2 and C2S2M2 PSII-LHCII supercomplexes placed in. Bottom panel: red outlines highlight possible in-plane clashes between different types of supercomplexes within the same membrane (outlines carried down from panels above). **B)** Quantification of the membrane coverage, depending on the type of supercomplexes placed in (the area of S2 LHCII trimers in supercomplexes is also shown). Reproduced results from the *C. reinhardtii* study (Chlamy) shown for comparison [2]. Error bars: SD. **C)** Cumulative membrane coverage of PSII supercomplexes throughout the stack. Each line in the plot represents the percentage of area occupied by PSII or different PSII-LHCII supercomplexes when adding membranes across the stack (from the first stacked membrane until the end of the granum). Line: mean, shading: SEM. Membranes from 15 grana were used for the analysis. **D)** Quantification of the in-plane clashes between PSII cores or different supercomplexes. PSII-PSII clash: 0.0002% (within error range); these might be complexes in direct contact like the example in A. **E)** Top: cartoon representation of the fifth membrane (M5) from the same granum with all C2S2 supercomplexes in place. Bottom, overlay of M4 (blue) and M5 (magenta) showing the overlap (white) between PSII-LHCII in appressed membranes separated by a stromal gap. **F)** Quantification of the stromal overlap between PSII cores, LHCII S2 trimers, and different supercomplex variants. For all plots, data was generated from 3 chloroplast preparations, using the best resolved membranes (n=173).

We do not observe instances where PSII complexes directly face each other across the luminal gap (Figure 8A-B, Figure 8 – video supplements 1 and 2). Rather, PSII densities interdigitate despite the thylakoid lumen being wide enough to accommodate two directly apposed complexes (Figure 4). Indeed, the luminal overlap is negligible between all visible densities (including cyt*b_6_f* and unknown particles) for the two membranes of an appressed grana thylakoid (e.g., M19-M20 and M21-M22 membrane pairs in Figure 8B). Similarly, across the stromal gap, the overlap of all densities is only slightly higher (∼4%) (e.g., M20-M21 membrane pair in Figure 8B). Performing this stromal overlap quantification with supercomplex footprints (Figure 7E) showed that PSII cores have 9% of their surface area overlapping, whereas C_2_S_2_ and C_2_S_2_M_2_ supercomplexes show 22% and 38% overlap of their area, respectively (Figure 7F). Within intact chloroplasts, we did not observe patterns of PSII supercomplexes aligning across appressed membranes (Figure 8A-B), contrary to the ordered lattices of PSII previously observed in thylakoids isolated from pea [28]. However, in tomograms of chloroplasts ruptured during blotting (e.g., Figure 1 – figure supplement 1F), we did observe that PSII densities sometimes align into semi-crystalline lattices with strong overlap between the PSII complexes from adjacent appressed membranes (Figure 8C-D). It is therefore plausible that extraction of thylakoids away from the chloroplast environment can induce the formation of PSII lattices between appressed membranes, some of which may have undergone restacking.

**Figure 8.**
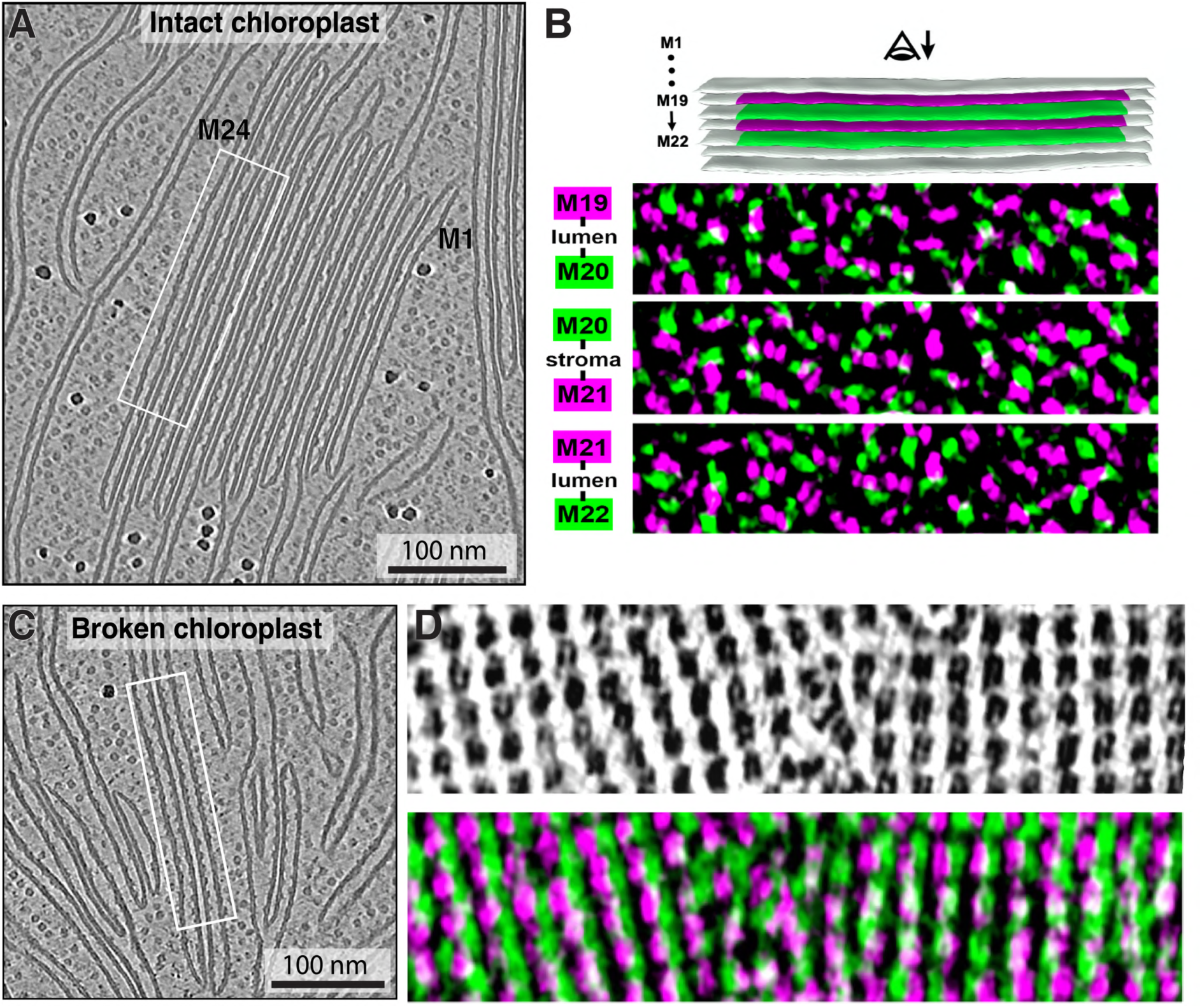
Overlap of densities across stacked membranes: intact vs. broken chloroplasts. the piece of a granum from A with four membranes highlighted (M19-M22). Odd numbered membranes in magenta, even numbered in green. Below: corresponding membranogram overlays of the adjacent membrane pairs. Coloring represents all EM densities protruding ∼2 nm from the luminal side of each membrane. The three examples show two luminal overlaps (top and bottom) and an overlap across a stromal gap (middle). White: overlap between EM densities. **C)** Tomographic slice showing a part of a broken chloroplast, containing individual membranes and potentially re-stacked thylakoids. White box marks the pieces of two membranes from adjacent thylakoids. **D)** Top: membranogram from a piece of one appressed membrane in C, showing densities protruding ∼2 nm from the membrane’s luminal surface. Note the arrays of PSII complexes and absence of any other type of particles. Bottom: membranograms from the pair of appressed membranes (green and magenta) overlayed with each other. White: overlap between EM densities. See Figure 8 – video supplements 1 and 2 for tomographic slices through the volumes shown in A and C.

## Conclusions and Speculation

Our cryo-ET study of intact spinach chloroplasts provides insights that span scales from the level of local membrane architecture (fine 3D ultrastructure of individual grana and connecting stromal lamellae) to the level of how individual protein complexes are organized within these membranes.

The architecture of spinach thylakoids visualized here is consistent with the previously proposed helical staircase models [24, 61, 62], although the structural details we observe are perhaps less stringent. The grana stacks of appressed membranes are not completely cylindrical: they have a wavy circumference (when viewed from the top, e.g., Figure 3), vary in diameter along the stack, and sometimes nearly merge with one another. We observe a variety of membrane architectures at the interconnections between appressed and non-appressed regions. Sometimes, the appressed domains shift smoothly into the non-appressed domains without architectural contortions (Figure 5, Figure 5 – figure supplement 1C), similar to what was observed in *C. reinhardtii*. However, much more complex transitions were also observed, with narrow membrane bridges or sharp forks (Figure 1 – figure supplement 2). At this level of membrane architecture, we also observed thylakoid contacts with plastoglobules (Figure 2) and the chloroplast envelope (Figure 1 – figure supplement 2). The pristine sample preservation inside vitrified chloroplasts, combined with the high resolution of our cryo-ET imaging, allowed us to measure thylakoid morphometrics (bilayer thickness, stromal and luminal gaps) with sub-nanometer precision (Figure 4).

At the level of molecular organization, the appressed and non-appressed membranes have markedly different composition, clearly visualized by the protruding protein densities in membranograms (Figure 5, Figure 5 – figure supplement 1). Similar to *C. reinhardtii* [2], there is a precise segregation between the two domains, with PSII restricted to the appressed membranes and PSI restricted to the non-appressed membranes. Cyt*b_6_f* is present in both domains. In contrast to previous biochemical studies, no obvious difference was seen between sub-regions of the non-appressed membranes (stromal lamellae, grana margins, grana end membranes) (Figure 5C, E; Figure 5 – figure supplement 1). Similarly, our analysis of PSII distribution in the appressed grana membranes showed that these complexes are uniformly distributed through the stack (Figure 6, Figure 7), with no changes in concentration observed across the grana layers nor close to the granum edge, where it meets the non-appressed membrane region. This confirms that, at least on the level of PSII distribution, there are no organizational differences across the granum.

Together, these observations support a simple two-domain model for thylakoid lateral heterogeneity *in vivo*, which states that 1) the appressed and non-appressed domains each have a homogeneous mix of their respective proteins throughout, and 2) there is an abrupt and discrete transition between the domains, with no visible mixing between the PSI and PSII complexes. Nonetheless, the domain boundary (grana margin) must be sufficiently fluid to facilitate the exchange of protein complexes during dynamic processes such as PSII repair and state transitions.

In the intact chloroplasts, we did not observe ordered lattices of PSII that were previously reported in less intact samples from various species [28, 48, 63, 64]. However, we did see semi-crystalline arrangements of PSII complexes in chloroplasts that had ruptured on the EM grid (Figure 8, Figure 8 – video supplements 1 and 2). Rearrangement of PSII into ordered arrays may therefore be triggered by exposure of the thylakoid network to a non-native environment with different buffering properties [65–67]. It remains to be determined whether PSII lattices could also have a physiological function *in vivo* [68] (to the best of our knowledge, the only *in situ* report of PSII arrays is in the dehydrated state of the resurrection plant *Craterostigma pumilum* [69]).

*In situ* cryo-ET only provides snapshots of living systems, but perhaps we can infer hints about the dynamic nature of thylakoids from these frozen moments in time. Our results show that PSII-LHCII supercomplexes do not align with each other under standard growth conditions. This distribution is consistent with supercomplexes mixing freely within the appressed domains without persistent static interactions with other supercomplexes. Based on evidence that LHCII plays an important role in thylakoid stacking [28, 45, 70–73], we hypothesize that while LHCII trimers can make relatively stable in-plane interactions to form PSII-LHCII supercomplexes, the other interactions within an appressed membrane and across the stromal gap to an adjacent stacked membrane are more transient. LHCII trimers are intrinsically multivalent and make electrostatic interactions with each other across the stromal gap using their disordered, positively-charged N-terminus [73]. Such a network of weak and multivalent interactions is reminiscent of the organizing principles of liquid-liquid phase separation [74, 75]. We speculate that stacked thylakoids in both green algae and plants are a condensation of membrane-embedded PSII-LHCII supercomplexes and free LHCII proteins. This condensation is driven by an ensemble of interactions both within the membrane plane and between stacked membranes (the latter via the N-terminus of LHCII), which are modulated by macromolecular crowding effects and membrane surface charge, respectively [67, 76, 77]. In contrast to canonical globular condensates, we propose that thylakoid stacks are multiple interacting layers of planar condensates. In addition to being a main driving force for the condensation of LHCII and PSII, these interactions between membrane layers also likely provide the mechanism for sterically excluding ATPsyn and PSI from the condensate, while permitting entry of cyt*b_6_f* because it lacks a large stromal density. An important prediction of this stacking condensation model is that PSII, LHCII, and cyt*b_6_f* are mobile within the appressed regions; however, we lack the methodology to directly track movements of single protein complexes within a native grana membrane.

## Limitations and Future Perspectives

Plant cells are generally difficult to vitrify and are too large for conventional FIB milling workflows. We opted for chloroplast extraction and plunge-freezing (Figure 1 – figure supplement 1), which facilitated the preparation, however not without disadvantages. The minutes-long isolation protocol can lead to relaxation of the physiological state of the chloroplasts, especially if it is temperature- or light-dependent. Moreover, plunge-freezing on EM grids can rupture the chloroplast envelope (Figure 1 – figure supplement 1B, F) and promote preferential orientation of the plastids — most often the entire thylakoid network lies flat on the surface of the grid support. Flat membranes, orthogonal to the electron beam during cryo-ET acquisition, provide top views (Figure 3, Figure 3 – figure supplement 1) that suffer from missing-wedge effects and are difficult to segment and analyze. We collected most of the tomograms towards the edges of the chloroplasts where this effect is less pronounced (more frequent side views), but this approach could induce region-specific bias.

Although more technically demanding, recent advances in cryo-FIB lift-out of high-pressure-frozen multicellular samples [78, 79] open future possibilities to perform cryo-ET on chloroplasts within native leaf tissue. Still, the relatively low throughput of cryo-ET and intrinsically small volume imaged by this technique present challenges for answering micron-scale questions about the overall membrane ultrastructure of the chloroplast; those questions are being successfully tackled by other microscopy techniques that image larger sample volumes at the trade-off of resolution (e.g., [22–24, 27, 61, 80, 81]). A more complete picture of thylakoid networks across length and time scales will require integrating the native molecular views provided by cryo-ET with complementary structural and cellular techniques [82]. Thylakoids are dynamic systems that adapt their molecular architecture to modulate their function in response to environmental changes. A future challenge is to accurately replicate this complexity with mesoscale modeling that incorporates parameters from diverse experiments to bridge thylakoid structure and function.

## Supporting information

Video supplements for Figures 1, 3 and 8

## Acknowledgements

We thank the Plant and Greenhouse facility of LMU Munich and especially Anja Schneider for help with the cultivation of plants. We thank Jürgen Plitzko and Wolfgang Baumeister at the Max Planck Institute of Biochemistry for access to electron microscopes. We thank Sebastian Ziegler and Fabian Isensee from the Applied Computer Vision Lab (ACVL) at the German Cancer Research Center (DKFZ) for their support in designing the membrane segmentation framework. L.L. also thanks Julia Schnabel from the Institute of Machine Learning in Biomedical Imaging (IML) at Helmholtz Munich for advice and supervision. Cryo-ET analysis was performed at the sciCORE (http://scicore.unibas.ch/) scientific computing center at the University of Basel. B.D.E. and W.W. acknowledge funding from a Human Frontier Science Program (HFSP) research grant (award number RGP0005/2021) and European Research Council (ERC) consolidator grant “cryOcean” (fulfilled by the Swiss State Secretariat for Education, Research and Innovation, award number M822.00045). M.P.J. was supported the grants from Biotechnology and Biological Sciences Research Council (BBSRC) (award number BB/V006630/1) and the Leverhulme Trust (award numbers RPG-2019-045 and RPG-2021-345). L.M. was supported by a BBSRC White Rose DTP studentship in Mechanistic Biology. L.L. acknowledges support from the Munich School for Data Science (MUDS) and a fellowship from the Boehringer Ingelheim Fonds.

## Data availability

The raw cryo-ET data, cryoCARE denoised tomograms and selected segmentation volumes are available at the Electron Microscopy Public Image Archive (EMPIAR), accession code: EMPIAR-12612. Positions of PSII particles used in the study are available in .star format at: 10.5281/zenodo.15090119. Denoised tomograms and segmentations shown in the figures are deposited in the Electron Microscopy Data Bank (EMDB), accession codes: EMDB-52542, EMDB-52543, EMDB-52544, EMDB-52545, EMDB-52546, EMDB-52547, EMDB-52548.

## Materials and Methods

### Plant growth

*Spinacia oleracea* WT plants were germinated and grown in a growth chamber for 6 weeks with a 12 h day/night cycle at 21-23°C and white-light illumination (150 µmol photons m^−2^s^−1^). One day before the experiment, the night cycle was extended to 20 h to minimize starch content in the chloroplasts. Prior to chloroplast isolation, plants were transferred from darkness to illumination for 1 h to acclimate them to light. This experiment was performed in three independent batches.

### Chloroplast isolation

Approximately 80 grams of fresh leaf mass was collected and immediately blended with 160 mL of isolation buffer (0.45 M Sorbitol, 20 mM Tricine-KOH, 10 mM EDTA, 10 mM NaHCO3, 5 mM MgCl2, 0.1% BSA, 0.2% D-Ascorbate) on ice. The slurry was filtered twice through cotton and a double layer of Miracloth to remove debris. The filtrate was then deposited on a 50% percoll cushion in isolation buffer and centrifuged for 8-10 mins at 7500 rpm. The green band in between percoll layers was collected and diluted in the isolation buffer at least 3 times (dilution depended on the efficiency of purification and type of the EM grids to be used). Chloroplast were plunge-frozen immediately afterwards.

### Plunge-freezing

Plunging was performed using a Vitrobot Mark 4 (Thermo Fisher Scientific). 4 µL of chloroplasts solution was placed onto R2/1 or R1.2/1.3 carbon-coated 200-mesh copper EM grids (Quantifoil Micro Tools), blotted for 7-8 seconds from the back side (front side pad covered with a Teflon sheet) with blot-force 10, and plunge-frozen in a liquid ethane/propane mixture cooled with liquid nitrogen. Grids were clipped into “autogrid” support rings modified with a cut-out on one side (FEI, Thermo Fisher Scientific) and stored in plastic boxes in liquid nitrogen until used for cryo-FIB milling.

### Focused Ion Beam milling

Cryo-FIB milling was performed as described previously [31] using an Aquilos dual-beam FIB/SEM instrument (Thermo Fisher Scientific). Grids were screened for good chloroplast coverage and coated with a layer of organometallic platinum using a gas injection system to protect the sample surface. The sample was milled with a gallium ion beam to produce ∼100-150 nm-thick lamellae. Some batches of lamellae were additionally sputter-coated with platinum at the end of the milling process. Grids were then transferred in liquid nitrogen to the TEM microscope for tomogram acquisition.

### Cryo-ET data acquisition

Tomographic data was collected on a 300 kV Titan Krios TEM instrument (Thermo Fisher Scientific), equipped with a post-column energy filter (Quantum, Gatan), and a direct detector camera (K2 Summit, Gatan). Tilt series were acquired using SerialEM software [83] with a dose symmetric scheme [84] starting at the pretilt of 10/-10° (depending on the loading direction of the grids), with 2° increment between tilts and spanning a total of 120°. Individual tilts were recorded in counting mode with an imaging rate of 8-12 frames per second, at a pixel size of 3.52 Å and target defocus in the range of 2.5-5 µm. The total accumulated dose for the tilt-series was kept at approximately 120 e-/Å^2^.

### Tomogram reconstruction and processing

All tilt-series were processed using the TOMOMAN pipeline [85]. Raw frames were aligned using MotionCor2 (v.1.4.0) [86]. Bad tilts were manually removed, and cleaned tilt-series were dose-weighted. The assembled tilt-series were aligned with patch tracking using the Etomo program from the IMOD (v.4.11.1) package [87, 88]. Final tomograms were reconstructed by weighted back projection in IMOD. To enhance contrast, the cryo-CARE denoising was applied [36]. For the views in Figure 3, we instead performed denoising with DeepDeWedge, which additionally attempts to fill in missing wedge information [37]. All images of tomograms (tomographic slices) shown were acquired using the IMOD 3dmod viewer.

### Membrane segmentation

Membrane segmentation was performed using AI-assisted approaches as described in [3]. Briefly, in order to generate membrane segmentations, we initially segmented the tomograms using the TomoSegMemTV segmentation protocol [89], and extracted patches of 160^3^ voxels of tomogram-segmentation pairs. After manually cleaning these segmented patches, we used these pairs to train a neural network (nnU-Net) to segment the entire thylakoid lattice. We predicted segmentations using the trained network, extracted and corrected new membrane patches, merged these into our training set, and started a new round of training. We followed this iterative approach until we were satisfied with the performance of our segmentation model. This model is now available in the MemBrain-seg package [3]. All segmentations shown in the figures were generated using this model, then cleaned in Amira (Thermo Fisher Scientific) and rendered in UCSF ChimeraX.

### Preparation of single-membrane segmentations

For all analyses requiring single membrane instances or stacks of individual membranes (e.g., particle organization in the membrane and overlays between membranes), the separate segmentations were made by cutting the full-tomogram segmentations from MemBrain-seg in Amira and exporting them as separate volume files (.mrc) and triangulated surface meshes.

### Measurement of appressed vs. non-appressed areas

For the quantification of appressed vs. non-appressed regions (shown in Figure 1E-F), we first segmented all membrane networks in all our tomograms using our trained MemBrain-seg model and then manually cleaned the output segmentations in Amira to remove tomogram edge artifacts. Afterwards, we converted the segmentations into triangular meshes using PyVista’s marching cubes algorithm [90], followed by Laplacian smoothing, giving normal vectors for each triangle on the mesh. For each triangle, we traced the distance to the next triangle along the normal vector. This next triangle along the normal belongs to potentially neighboring membranes, and by thresholding the distance, we divided the meshes into “appressed” and “non-appressed” triangles. To account for noisy distance predictions, we applied a majority vote among geodesically neighboring triangles to receive a smooth representation of appressed and non-appressed regions. Furthermore, by computing the principal curvature at each triangle, we excluded the membrane segmentation edges (e.g., clipped by the tomogram volume or ending due to mis-segmentation), as these have normal vectors pointing away from the stack and therefore would distort the metrics towards more non-appressed regions. By summing up the triangle areas of all remaining mesh triangles for both classes, we received the total areas of appressed and non-appressed membranes.

### Thylakoid spacing and thickness calculations

In order to measure distances between different membrane layers, stromal and luminal gaps, as well as entire thylakoid thickness (shown in Figure 4), we utilized membrane flattening functionalities from blik [91]. The goal of this step was to receive a “stack” of 2D images for each membrane, where each 2D image in the z-stack corresponds to the flattened front view of the membrane at a different distance along the membrane normal vector. This is to remove the effects of membrane curvature and increase signal-to-noise ratio of the measurements as well as to ensure sampling orthogonal to the plane of lipid bilayers. To perform the flattening, we first sampled a grid of points on each 3D membrane segmentation. This grid allowed blik to generate the 2D image stacks along the normal vectors. Next, we divided each membrane into 50 nm patches. By averaging the intensities of each 2D image in these stacks, we received a single value for each z-coordinate, leading to a 1D plot along the z-axis, i.e., the membrane normal vector. From this 1D plot, we extracted local intensity minima and maxima corresponding to center points of lipid bilayers. After manual cleaning of outlier plots (i.e., plots where the automated assignment of keypoints failed), this representation allowed us to estimate the distances between different membranes and gaps. To account for the soft edge of the membrane in cryo-ET data, we selected a half-way point between the maximal signal of the membrane leaflet to the center of the membrane as the “edge” and applied this correction symmetrically on both sides of the signal peak for more accurate measurements. See Figure 4 - figure supplement 1 for the visual description.

### Membrane protein ground truth positions

For training MemBrain, our membrane protein localization network [3], we required ground truth positions for the PSII proteins. To generate them, we imported the single membrane .obj files into Membranorama (github.com/dtegunov/membranorama) to visualize the membranes and their embedded proteins by projecting the tomogram densities onto the surface mesh. In this program, we were able to manually annotate PSII positions and orientations in 45 membrane patches from seven different tomograms, which we then used as training data for the MemBrain model.

### MemBrain training

We utilized our membrane protein ground truth data to train a MemBrain model. To do this, we split our full dataset consisting of 45 membranes into a training set (28 membranes), a validation set (7 membranes), and a test set (10 membranes). As a required first step in the MemBrain workflow, we manually defined the side of each membrane facing toward the thylakoid lumen as the side of interest. Then, we trained MemBrain using its default parameters to predict “heatmaps” depicting each membrane point’s distance to the closest PSII. Using our ground truth annotations (validation set), we further tuned the best bandwidth parameter for the subsequent Mean Shift clustering, which we determined to be 21 bin4 voxels, corresponding to 29.6 nm

### Application of trained MemBrain model

After training and tuning MemBrain, we applied it to the full dataset, consisting of 455 discrete membrane segmentations from the nine best quality tomograms. This gave us good initial predictions for positions and orientations of PSII complexes in all membranes. In order to confirm their validity, we inspected all 455 membranes again, discarded 152 membranes due to insufficient quality, and manually corrected false MemBrain predictions. The remaining 303 membranes were used for the analysis. Wherever image quality allowed, we also annotated cyt*b_6_f* positions. This gave us an accurate membrane protein dataset, consisting of 14433 PSII, 937 cyt*b_6_f*, and 5934 positions of unknown particles (non-identifiable densities).

### Center-to-center distances

In order to analyze the nearest neighbor distances between our particle positions, we first converted each voxel-wise membrane segmentation into a surface mesh representation using PyVista’s marching cubes algorithm, followed by Laplacian surface smoothing (applied to individual single membrane instances). This allowed us to compute exact geodesic distances between two positions on the mesh using The Virtual Brain’s (https://github.com/the-virtual-brain) implementation of the discrete geodesic distance algorithm [92]. For each particle location, we computed the geodesic distance to the center position of the nearest neighbor from the respective target particle class (e.g., PSII-NN_Spin_, cyt*b_6_f*--NN_Spin_, all-NN_Spin_; shown in Figure 6B).

### Distance to appressed region edge

To analyze the change of protein distribution with respect to distance from the edge of appressed membrane regions, we first annotated these membrane edges manually using Membranorama. We then divided the Euclidean distance to the membrane edge into 10nm-sized bins and computed the total area of mesh triangles within each bin. By dividing the number of particle positions by the triangle area within each bin, we obtain average protein complex concentrations per bin. Analogously, for each bin, we computed the average nearest neighbor distances. Plotting these concentrations against the respective bin distances provided the graphs displayed in Figure 6D.

### Overlap analysis

To analyze the overlap between PSII-LHCII supercomplexes in neighboring membranes (Figure 7E and F) or cumulative occupancy in membranes across the stack (Figure 7C), we projected protein positions and orientations onto a common membrane surface. We calculated their overlaps by comparing the overlaying triangles on the segmented membrane. To process a stack of membranes, we calculated a connection vector between the first and last membrane of the stack by computing the average membrane normal vector for the starting membrane. Then, for each membrane, we projected the corresponding membrane protein positions along the normal vector onto the starting membrane mesh. Mapping the protein complex footprint (based on its 3D structure filtered to 14 Å resolution) into the correct orientation in this new position allowed us to calculate the area on the membrane surface that was occupied by the respective complexes. Doing this for all membranes gave us a set of occupied areas per membrane, which we then analyzed for overlapping areas. To avoid bias due to membrane misalignment (e.g., stack being inclined in the tomogram slab resulting in incomplete overlap between membranes), we also projected the segmentations of all membranes onto the starting mesh and only analyzed the areas that were shared by all projected membrane segmentations.

## Supplemental Figures

**Figure 1 – figure supplement 1.**
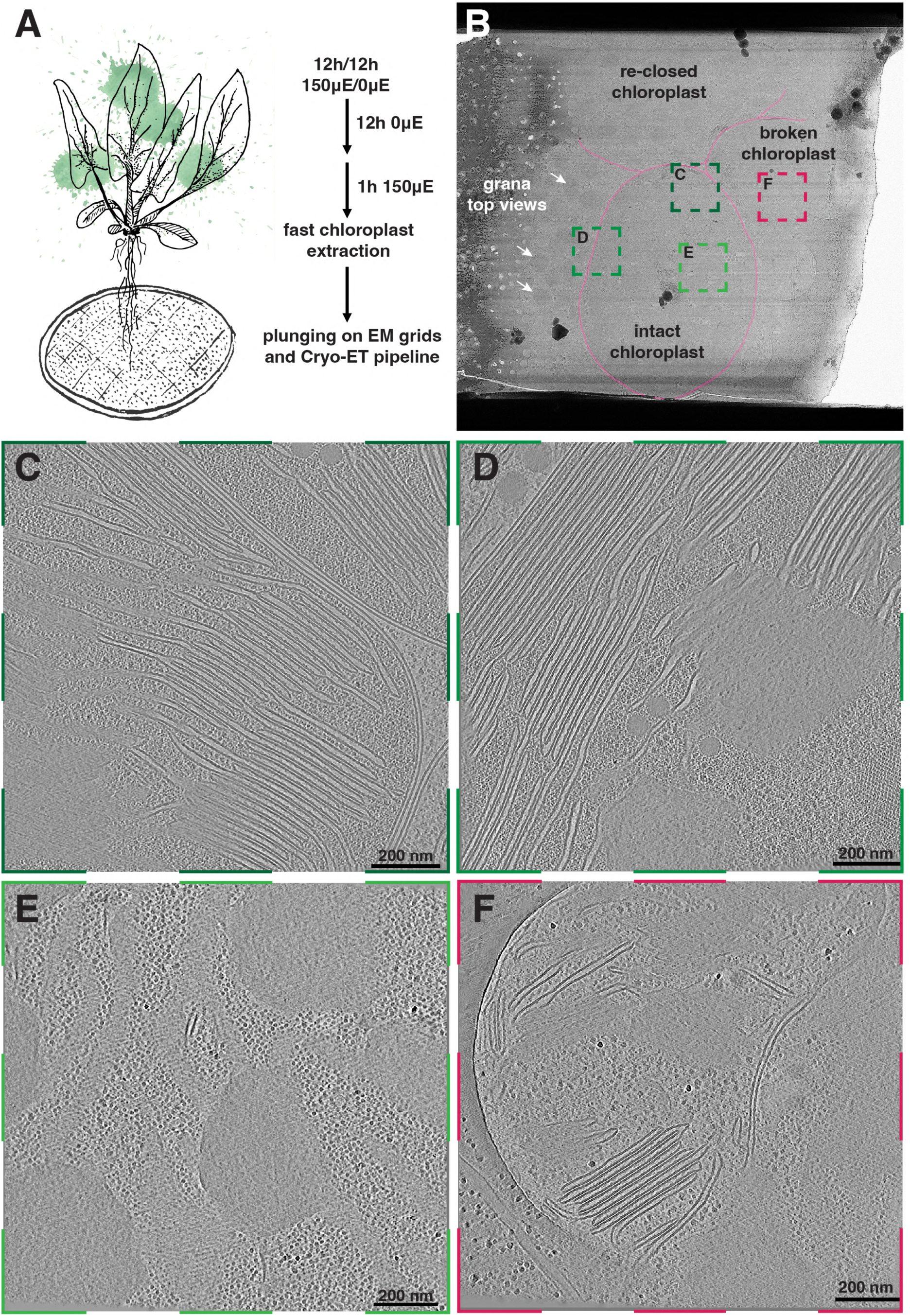
Sample preparation, tomogram selection on the FIB-milled lamellae, and types of tomograms collected. **A)** Simplified schematic of the chloroplast preparation protocol, which aimed to minimize the time between chloroplast isolation and freezing. **B)** An overview of a typical FIB-milled lamella containing chloroplasts as seen in the TEM microscope. Chloroplasts can break and sometimes re-close during the isolation and plunging procedures. We tried to identify intact, spheroidal chloroplast sections in the lamellae and focused tomogram collection in those regions. White arrows point the grana stacks, visible in the overviews. **C-F)** Examples of tomograms collected in different regions of an intact chloroplast (C-E) as well as in a region containing a broken chloroplast (F).

**Figure 1 – figure supplement 2.**
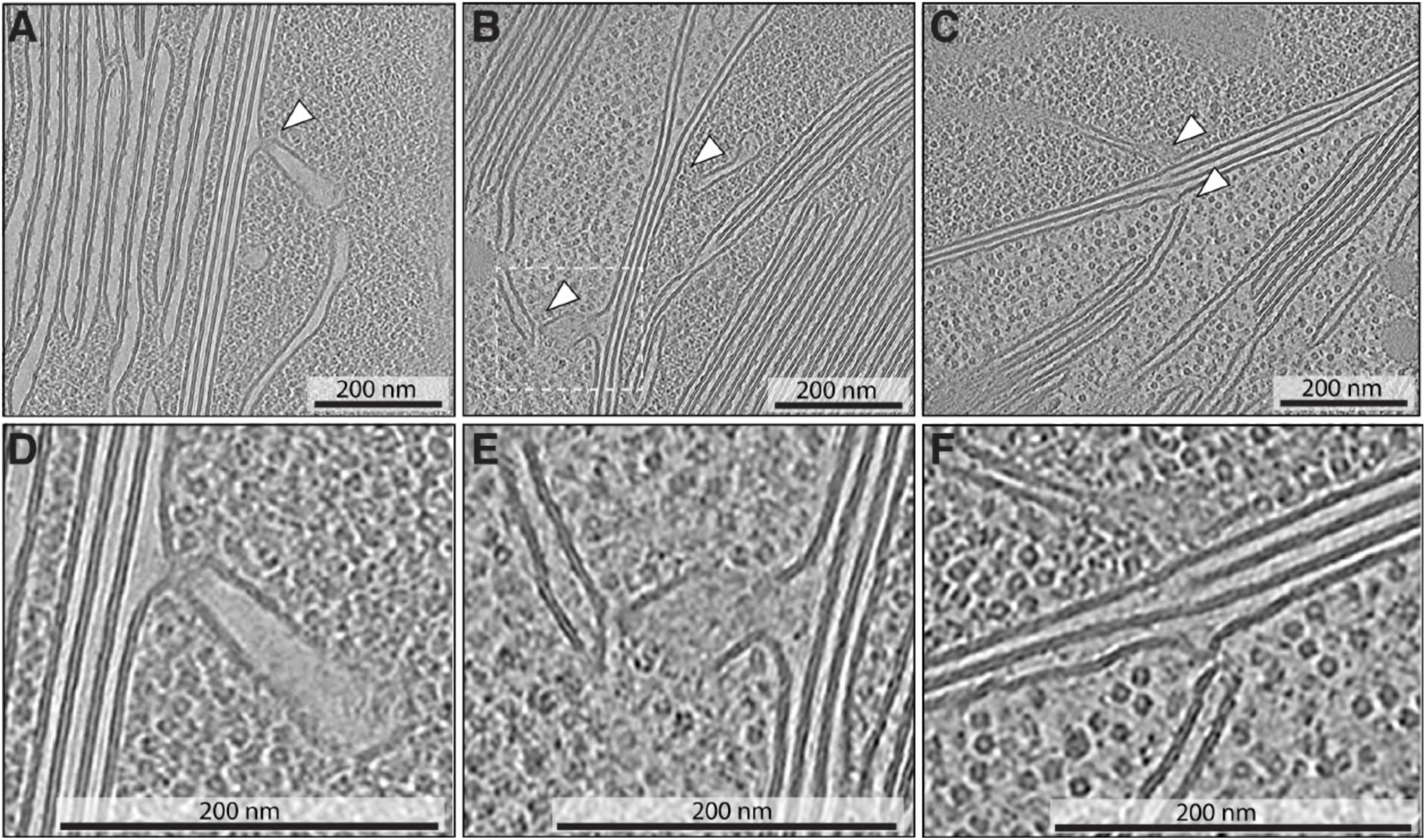
Thylakoid-chloroplast envelope contacts. Rare instances of thylakoids and thylakoid-like membranes approaching and contacting the inner envelope membrane inside intact chloroplasts. **A-C)** Tomographic slices showing overviews of the contact sites (white arrowheads). **D-F)** Zoom-ins of the contacts from respective images. Dashed box in B corresponds to the enlarged view in E.

**Figure 1 – figure supplement 3.**
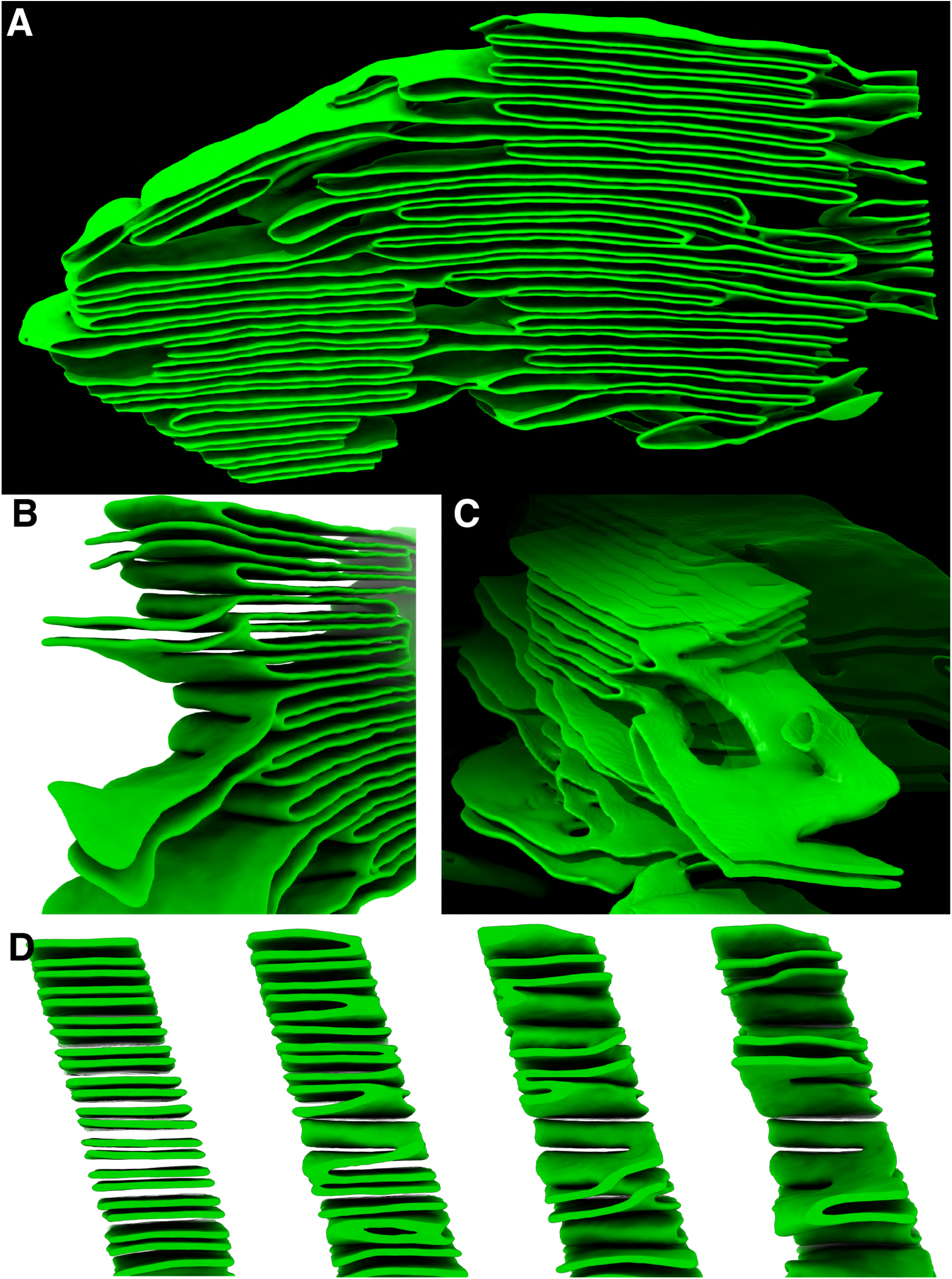
Ultrastructure of grana sides and transition to stromal lamellae. **A)** Segmentation of thylakoid membranes from two grana in close proximity. Note the irregular stacks and shifted thylakoids. **B)** View of the margin region on one side. Grana-forming thylakoids are flat and straight, whereas stromal lamellae are more undulating. **C)** View of the other side. Note the convoluted shape of stromal lamellae connecting with two other lamellae spanning from the same granum. The top part of the membrane segmentation was set to high transparency to improve visibility. **D)** Serial perpendicular slices through the granum, showing appressed membranes transitioning into stromal lamellae. Membranes merge and form bridges between each other.

**Figure 3 – figure supplement 1.**
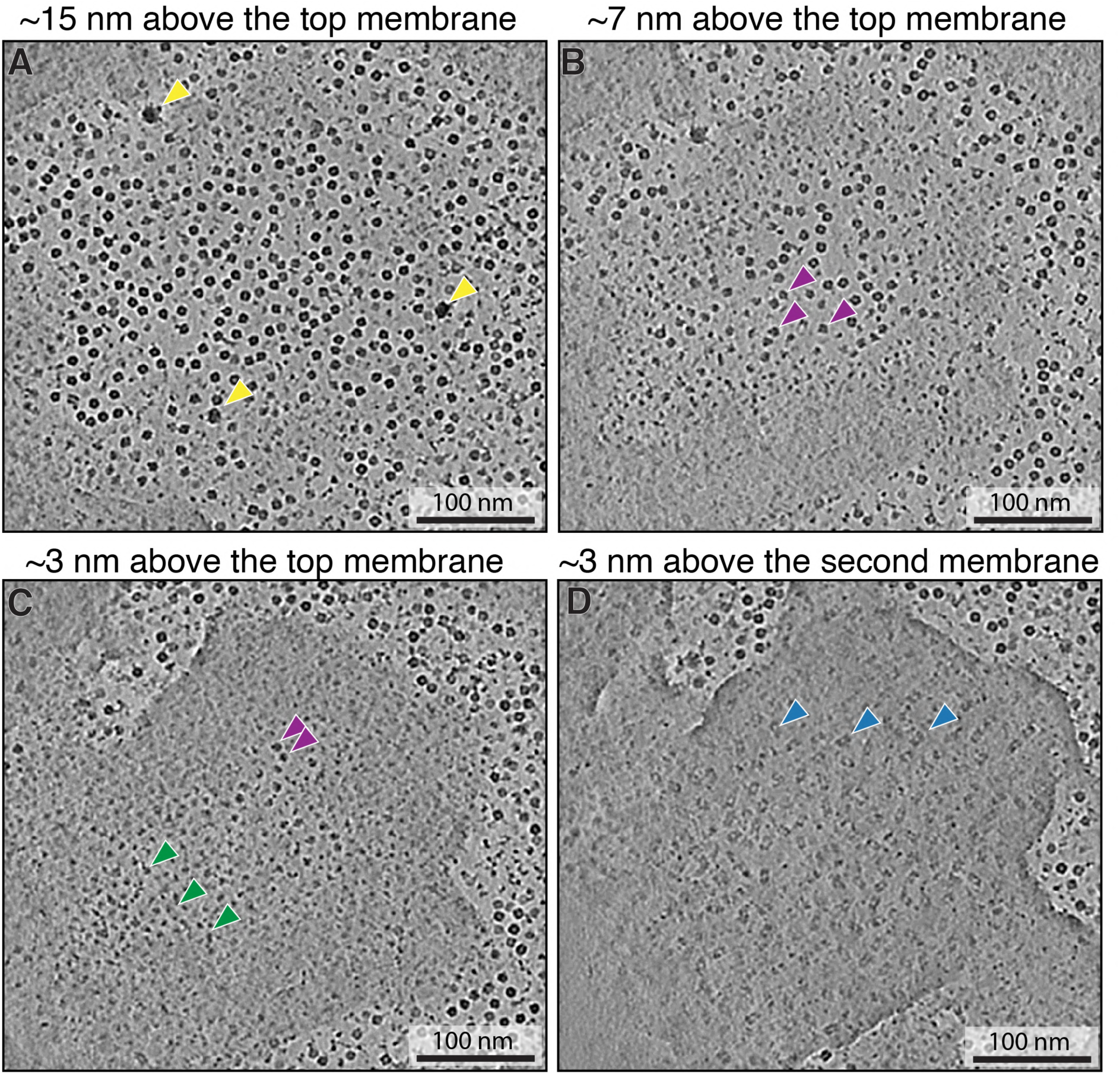
Top view of a granum. Tomographic slices from a tomogram denoised with DeepDeWedge program [37] (high contrast, missing wedge inpainting) showing the chloroplast stroma 15 nm above the granum (**A**), 7 nm above the granum (**B**), and 3 nm above the top membrane (**C**), as well as the luminal space between the top and second membranes of the granum (**D**). Note the high number of globular particles (mostly Rubisco) in A. Arrowheads indicate complexes appearing in the slices (Yellow: ribosomes, purple: ATPsyn, Green: putative PSI, Blue: PSII). Note the similar size of Rubisco and the ATPsyn F1 subunit, which complicates assignment. The tomographic volume was rotated so the top membrane was approximately parallel to the xy-plane.

**Figure 4 – figure supplement 1.**
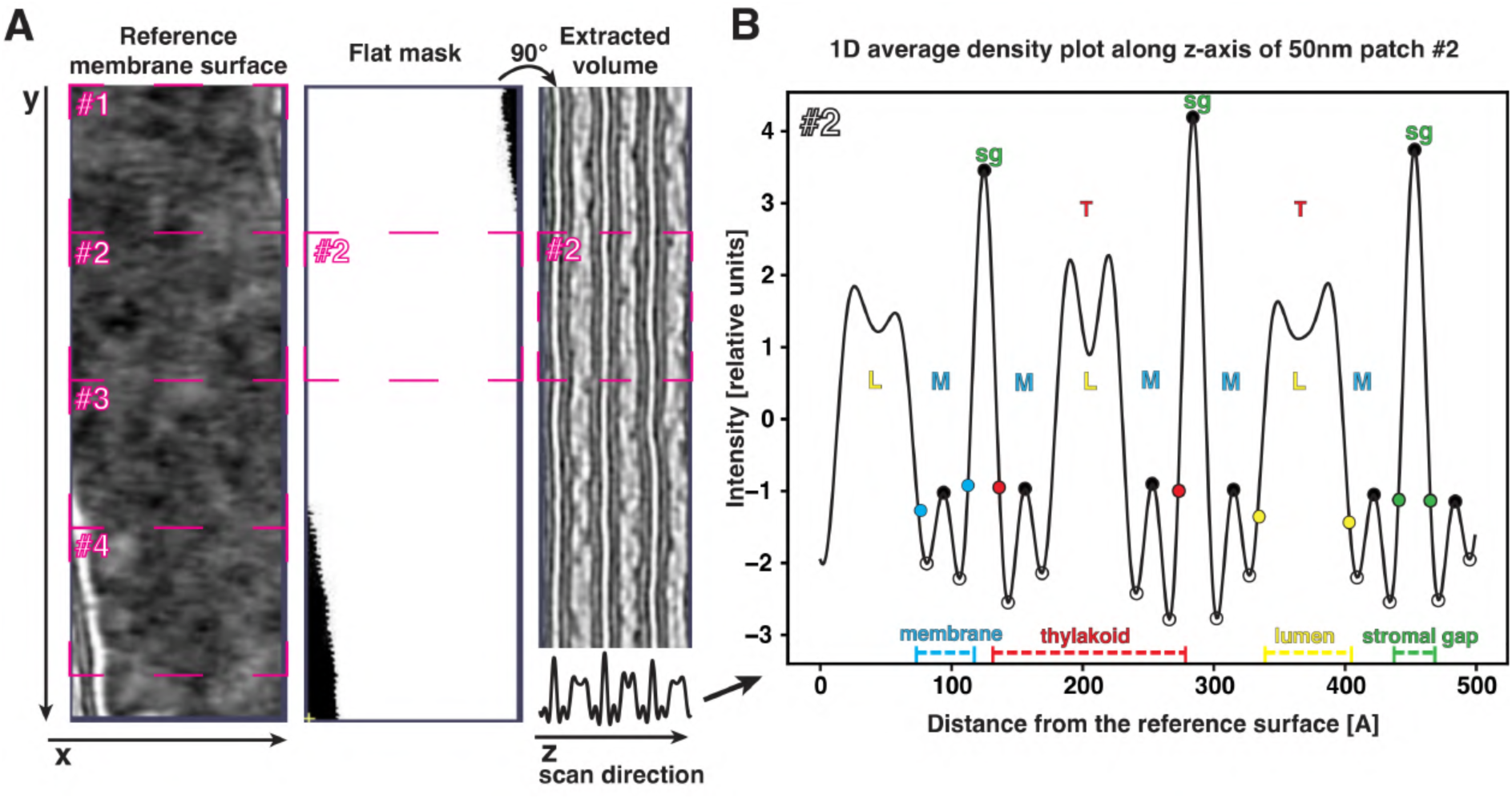
Methodology for measuring thylakoid widths from tomograms. **A)** Flattened and aligned thylakoid volume (processed in blik [91]) extracted around a representative membrane instance. Left: Top view of the flat stack at center z-value (middle of a membrane bilayer). The membrane is divided into 4 patches of 50 nm length along the y-axis (pink rectangles and numbers). Middle: Membrane mask corresponding to the z-slice to focus analysis on membrane regions. Right: x-z view of the flattened volume. Note the parallel orientation of the neighboring membranes. Density scan is performed by averaging all values for each z-value in the volume (here, patch #2), corresponding to averaging along the membrane normal vector. Density values are plotted as a 1D line profile (bottom part of the extracted volume panel shows the simplified plot; note the oscillation of the intensity signal across membrane stack). **B)** 1D density profile of patch #2 shown in A. Local minima and maxima are indicated with black and white dots, respectively. Blue, red, yellow, and green dots represent approximated membrane edge positions. To approximate membrane edges, we first calculated the distance between bilayer minima (white dot) and center between bilayers (neighboring black dot). Then, we added half this distance to the bilayer minimum in the opposite direction. This approximation assumes that bilayer intensities fade out symmetrically on both sides, and sets an arbitrary, but consistent threshold. We calculated widths of different components: membranes (M, in blue), thylakoid (T, in red), lumen (L, in yellow) and stromal gap (sg, in green). These measurements were conducted for each patch from all available membranes. Summarized results from all measurements are presented in Figure 4.

**Figure 5 – figure supplement 1.**
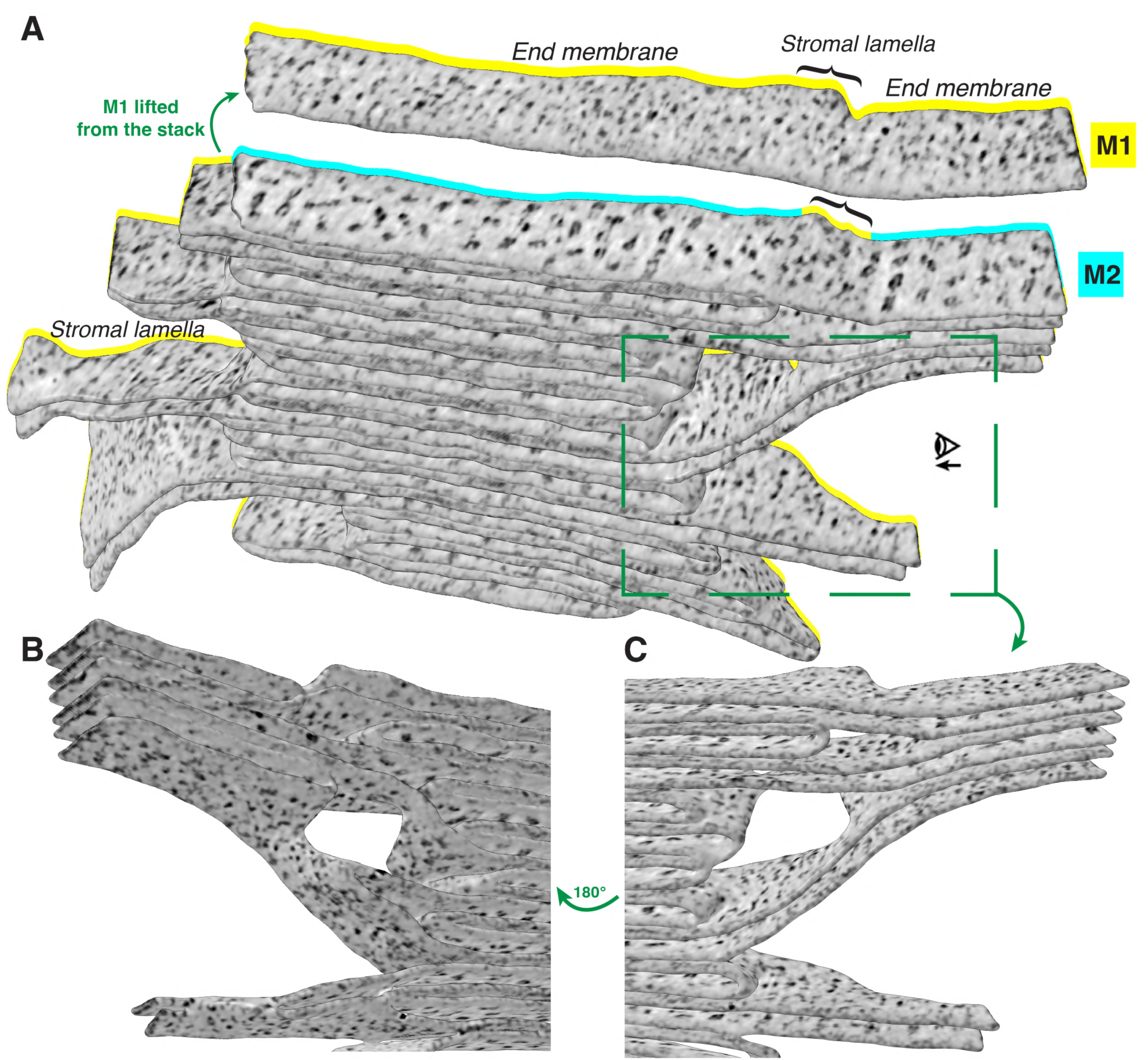
Protein densities on two domains of thylakoids. **A)** A 3D rendering of a thylakoid granum and stromal lamellae with tomographic densities projected onto the membrane surfaces (membranograms). The first membrane of the granum (M1) is separated from the stack to show the PSII and cyt.*b6f* densities in the appressed region of M2 below. Note that all non-appressed membranes (including grana margins, end membranes, and stromal lamellae) are populated with similar densities. **B)** and **C)** Zoom-in views of stromal lamellae extending from the side of the granum. The eye icon shows the viewing direction in C. All panels show membranes at the same threshold value and the same color scale.

## Supplemental Videos

**Figure 1 – video supplement 1.** Overview of a chloroplast tomogram, slicing through the tomographic volume and then revealing segmentations of the thylakoid and chloroplast envelope membranes.

**Figure 3 – video supplement 1.** Slices through the tomogram shown in Figure 3, showing the chloroplast stroma and top views of the thylakoid network, with stromal lamellae connecting the grana.

**Figure 8 – video supplement 1.** Slices through the tomogram shown in Figure 8A, highlighting the organization of PSII particles in near-native thylakoid membranes within an intact chloroplast.

**Figure 8 – video supplement 2.** Slices through the tomogram shown in Figure 8C, highlighting the organization of PSII particles in thylakoid membranes from a broken chloroplast

